# NF90 Modulates Processing of a Subset of Human Pri-miRNAs

**DOI:** 10.1101/2020.01.24.916957

**Authors:** Giuseppa Grasso, Takuma Higuchi, Jérôme Barbier, Marion Helsmoortel, Claudio Lorenzi, Gabriel Sanchez, Maxime Bello, William Ritchie, Shuji Sakamoto, Rosemary Kiernan

## Abstract

MicroRNAs are predicted to regulate the expression of more than 60% of mammalian genes and play fundamental roles in most biological processes. Deregulation of miRNA expression is a hallmark of most cancers and further investigation of mechanisms controlling miRNA biogenesis is needed. The dsRNA-binding protein, NF90 has been shown to act as a competitor of Microprocessor for a limited number of pri-miRNAs. Here, we show that NF90 has a more widespread effect on pri-miRNA biogenesis than previously thought. Genome-wide approaches revealed that NF90 is associated with the stem region of 38 pri-miRNAs, in a manner that is largely exclusive of Microprocessor. Following loss of NF90, 22 NF90-bound pri-miRNAs showed increased abundance of mature miRNA products. NF90-targeted pri-miRNAs are highly stable, having a lower free energy and fewer mismatches compared to all pri-miRNAs. Mutations leading to less stable structures reduced NF90 binding while increasing pri-miRNA stability led to acquisition of NF90 association, as determined by RNA EMSA. NF90-bound and modulated pri-miRNAs are embedded in introns of host genes and expression of several host genes is concomitantly modulated. These data suggest that NF90 controls the processing of a subset of highly stable, intronic miRNAs.

## INTRODUCTION

MicroRNAs (miRNAs) are short non-coding RNAs that negatively regulate the expression of a large proportion of cellular mRNAs, thus affecting a multitude of cellular and developmental pathways (1,2). The canonical miRNA biogenesis pathway involves two sequential processing events catalysed by RNase III enzymes. In the nucleus, the microprocessor complex, comprising the RNase III enzyme Drosha, the double-stranded RNA-binding protein, DGCR8 and additional proteins carries out the first processing event, which results in the production of precursor miRNAs (pre-miRNAs) (3,4). These are exported to the cytoplasm, where a second processing event carried out by another RNase III enzyme, Dicer, leads to the production of mature miRNAs that are loaded into the RISC complex (5).

Due to the central role of miRNAs in the control of gene expression, their levels must be tightly controlled. Indeed, deregulation of miRNA expression is associated with aberrant gene expression and leads to human disease (6–9). Consequently, miRNA biogenesis is tightly regulated at multiple steps, both transcriptional and post-transcriptional. Increasing evidence suggests that RNA binding proteins (RBPs) act as post-transcriptional regulators of miRNA processing. Many RBPs modulate the processing efficiency of Microprocessor, either positively or negatively, by binding to regions of the pri-miRNA. A number of RBPs have been shown to bind the terminal loop, which can either facilitate or inhibit cropping by Microprocessor. For example, LIN28B binds the terminal loop of pri-Let-7, which prevents its processing by Microprocessor (10). Binding of hnRNP A1 to the terminal loop has been shown to exert either positive or negative effects on Microprocessor activity, depending on the pri-miRNA target. It promotes cropping of pri-miR-18A while it inhibits processing of pri-let-7. KSRP is another terminal loop-binding RBP that facilitates Microprocessor cleavage of several pri-miRNA targets, including pri-let-7 where it acts as a competitor of hnRNP A1 (11,12). Several other RBPs, including SMAD, TPD-43, SRSF1 and RBFOX, have been shown to bind pri-miRNA terminal loops to influence Microprocessor activity (see (13) for review). In most cases, they have been shown to bind specific pri-miRNAs, such as pri-let-7, or a limited subset of pri-miRNAs. To date, only NF90/NF45 heterodimer and ADAR1,2 have been shown to bind the double stranded stem region of pri-miRNAs (14–17). Both factors negatively affect Microprocessor activity. Indeed, NF90 has been shown to bind double stranded RNA in a mode similar to that of ADAR2 (18). Like terminal loop binding RBPs, binding of NF90/NF45 or ADAR1,2 has thus far been demonstrated for a very limited number of pri-miRNAs. NF90 has been shown to associate with pri-miR-7-1, pri-let-7A and pri-miR-3173 in human cells (14,15,19).

We have previously shown that NF90 associates with pri-miR-3173, which is located in the first intron of Dicer pre-mRNA. Binding of NF90 prevented cropping of pri-miR-3173 by Microprocessor and promoted splicing of the intron, thereby facilitating expression of DICER. By modulating DICER expression, NF90 was found to be an independent prognostic marker of ovarian carcinoma progression (19). Levels of NF90 are known to be elevated in hepatocellular carcinoma (HCC) and the effect of NF90 on processing of pri-miR-7-1 contributes to cellular proliferation in HCC models (14,20). Here, we have used genome-wide approaches to identify pri-miRNAs that are associated with and modulated by NF90 in HepG2 model of HCC. We identified 38 pri-miRNAs that are associated with NF90, in a manner that is for the most part exclusive of Microprocessor. Of these, 22 showed increased abundance of mature miRNAs products upon loss of NF90. NF90-targeted pri-miRNAs appear to be highly stable, having a lower free energy and fewer mismatches compared to all pri-miRNAs. Destabilization of the structures by mutation reduced NF90 association as determined by RNA EMSA. Of the 22 NF90-modulated pri-miRNAs, 20 are embedded exclusively in introns of host genes. Transcriptomic analysis revealed that the expression of the host gene is concomitantly modulated for several, including an oncogene implicated in metastasis of hepatocellular carcinoma, TIAM2. These data suggest that NF90 controls the processing of a subset of intronic miRNAs, which in some cases affects the expression of the host gene.

## MATERIAL AND METHODS

### Cell culture

Human HepG2 cell line was grown in Dulbecco’s Modified Eagle’s medium – high glucose (Sigma-Aldrich®, D6429) supplemented with 10% fetal bovine serum (PAN Biotech, 8500-P131704), 1% penicillin-streptomicin (v/v) (Sigma Aldrich®, P4333) and 1% L-glutamine (v/v) (Sigma Aldrich®, G7513). Human HEK-293T cells were grown in Dulbecco’s Modified Eagle’s high glucose medium with HEPES (Sigma-Aldrich®, D6171) supplemented with 10% fetal bovine serum, 1% penicillin-streptomicin and 1% L-glutamine. Cells were cultured at 37°C in a humidified atmosphere containing 5% CO2. HepG2 were seeded at 1.5 × 10^6^ cells in 6-well plates the day of siRNA transfection or at 7.5 × 10^5^ cells in 6-well plates the day prior to plasmid transfection. HEK-293T were seeded at 6 × 10^5^ cells in 6-well plates the day of siRNA transfection.

### Transfection of small interfering RNAs

Double-stranded RNA oligonucleotides used for RNAi were purchased from Eurofins MWG Operon or Integrated DNA Technologies. Sequences of small interfering RNAs (siRNAs) used in this study have been described previously (19) and are shown in Supplementary Table S1.

HepG2 or HEK-293T cells were transfected with siRNA (30 nM final concentration) using INTERFERin^®^ siRNA transfection reagent (Polyplus Transfection) according to the manufacturer’s instructions. Two rounds of transfection were performed. The first transfection was carried out the day of seeding; on the fourth day cells were passaged and a second round of transfection was performed. Cells were collected for RNA extraction or protein purification approximately 65 h after the second transfection.

### Immunoblot

HepG2 were lysed using RIPA buffer (50 mM Tris-HCl pH=7.5, 150 mM NaCl, 1 % NP40, 0.5 % Sodium Deoxycholate, 0.1 % SDS, Halt^™^ Phosphatase Inhibitor Cocktail (Thermo Fisher Scientific)). Protein extracts (30 μg for NDUFS8, 50 μg for TIAM2 and 5 μg for all other proteins) were immunoblotted using the indicated primary antibodies (Table S2) and anti-mouse, anti-rabbit or antirat IgG-linked HRP secondary antibodies (GE Healthcare) followed by ECL (Advansta).

### Small RNA-seq and RNA-seq

Total RNA was extracted using TRIzol (Thermo Fisher Scientific) according to the manufacturer’s instructions. Small RNA-seq (single end, 50 bp) was carried out by BGI Genomic Services (HepG2) or Fasteris (HEK-293T) in triplicate samples. Raw data were processed using the Subread package (version 1.6.0) as previously described (21) and the reference annotation was obtained from miRBase release 22.1 database (22). Statistical analysis was performed using DESeq2 (version 2.11.40.2). RNA-seq (paired-end, 125 bp) was carried out by BGI Genomic Services in triplicates. Raw data were processed using HISAT2 (version 2.1.0) and featureCounts (version 1.6.3), statistical analysis was performed using DESeq2. Reference annotation was obtained from ENSEMBL (GRCh38.96).

### RT-qPCR, Modified 5’ RLM RACE and RNA EMSA

Total RNA was extracted from HepG2 cells using TRIzol reagent (Thermo Fisher Scientific) and RNA was treated with DNAse I (Promega) according to the manufacturer’s instructions. RNA was used for RT-PCR and modified 5’ RLM-RACE as described previously (19).

For RT-qPCR, RT was performed using TaqMan^™^ Reverse Transcription Reagent or TaqMan^™^ Advanced miRNA cDNA Synthesis Kit (Thermo Fisher). qPCRs were performed using GoTaq^®^ Probe qPCR Master Mix (Promega) or TaqMan^®^ Fast Advanced Master Mix (Thermo Fisher).

Modified 5’ RLM RACE was performed according to the manufacturer’s instructions (FirstChoice^™^ RLM-RACE kit, ThermoFisher Scientific). In order to detect premature miRNAs, the step using Calf Intestine Alkaline Phosphatase was omitted. Sequences of the primers used for PCR amplification are shown in Supplementary Table S3.

RNA EMSA was performed as described previously (15) using recombinant NF90. The pri-miRNA probes were amplified by PCR using the primers shown in Supplementary Table S3. Sequences of mutant pri-miRNAs are shown in Supplementary Table S4.

### Bioinformatic analyses

For splicing analysis, exon-intron junction reads and exon-exon junction reads were counted using IRFinder software (23), version 1.2.5. The number of exon-exon junction reads was divided by the number of exon-intron junction reads to obtain a spliced/unspliced ratio.

Enhanced UV crosslinking followed by immunoprecipitation (eCLIP) data for NF90, DGCR8 and DROSHA obtained in HepG2 cells by Nussbacher and Yeo (24) were retrieved from the NCBI database (NF90 eCLIP: ENCSR786TSC; DGCR8 eCLIP: ENCSR061SZV; DROSHA eCLIP: ENCSR834YLD). Peaks were filtered based on Fold Change (FC ≥ 1.5) and p-value (Bonferroni-Adj P-val ≤ 0.05). Distribution of eCLIP reads along the miRNAs was evaluated using deeptools software (version 3.1.3). Bigwig files from different replicates were merged using bigWigMerge v2. The base pair probability at each position of miRNA hairpins was calculated using RNAfold software (version 2.4.7).

Free energy analysis was performed using RNAfold software, version 2.4.7. Statistical analysis was performed using R (version 3.5.1).

Validated targets of the double positive miRNAs were extracted from MirTarBase database, release 7.0 (25). Gene ontology was performed on the expressed validated target using DAVID Functional Annotation Tool database version 6.8 (https://david.ncifcrf.gov) (26). Motif search was performed using MEME (version 5.0.5).

## RESULTS

### NF90 modulates the abundance of a subset of human miRNAs

To determine the effect of NF90 on the abundance of miRNAs, we performed small RNA-seq of biological triplicate samples obtained from HepG2 cells that had been transfected with a non-targeting control siRNA (siScr) or an siRNA targeting NF90 (siNF90) (Figure 1A, top panel). Of 1917 miRNA precursors annotated in miRBase, 1105, which corresponds to 1661 mature 5p and 3p miRNA products, were found to be expressed in HepG2 cells. Following loss of NF90, differential expression analysis (fold change ≥ 1.5 or ≤ 0.667; *AdjP-value* ≤ 0.05) showed that 212 precursor miRNAs, corresponding to 268 mature miRNAs, were upregulated while 126, corresponding to 149 mature miRNAs, were downregulated (Figure 1B). MiRNAs that have previously been shown to be modulated by NF90, miR-7-1 (14) and miR3173 (19), were found to be upregulated in HepG2 cells following loss of NF90 (Figure 1B, red dots).

**Figure 1.**
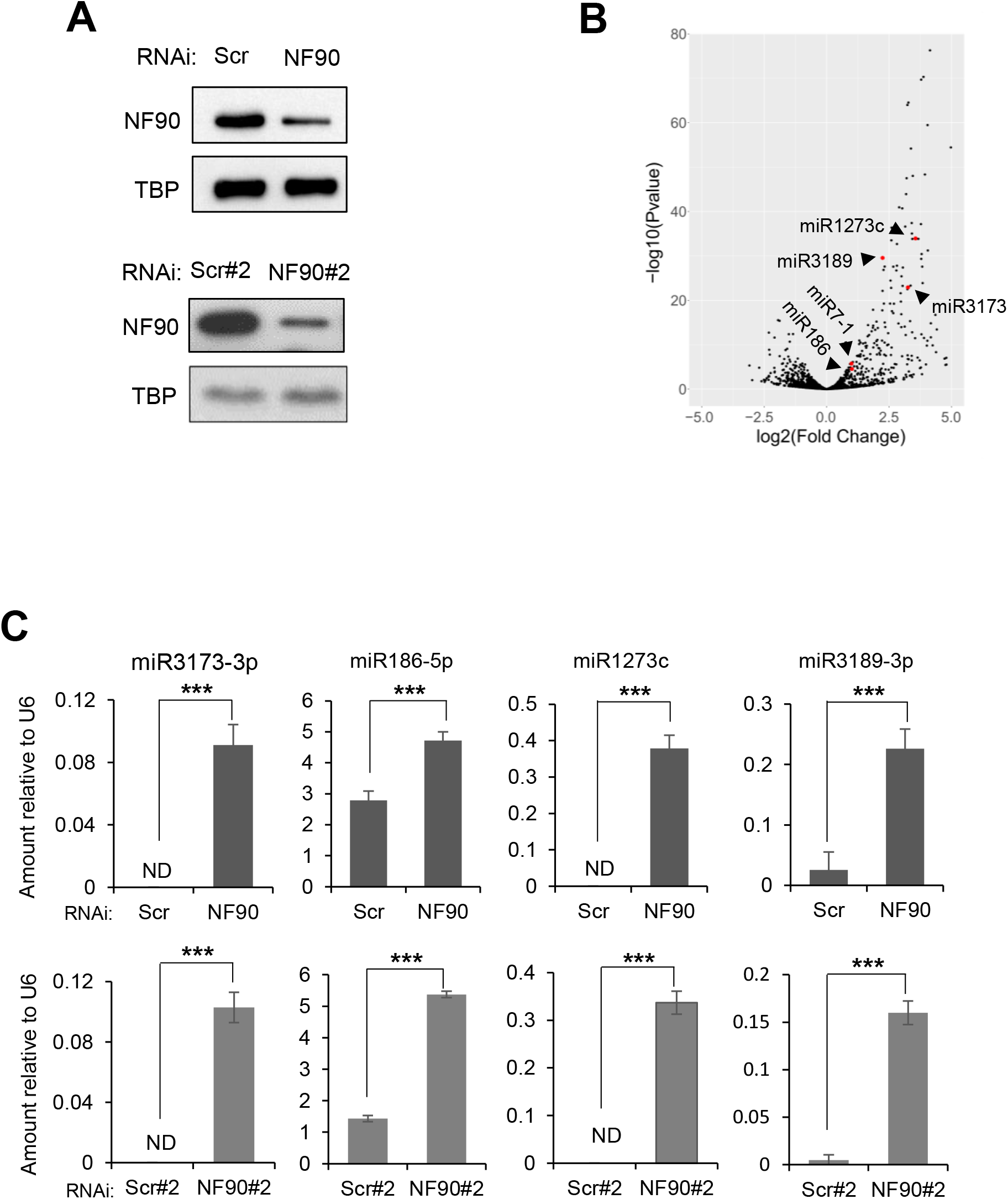
NF90 Modulates the Expression Level of a subset of miRNAs in HepG2 cells. (A) Extracts of HepG2 cells transfected with non-targeting control siRNAs (Scr, Scr#2) or siRNA targeting NF90 (NF90, NF90#2) as indicated were analyzed by immunoblot using the antibodies indicated. (B) Total RNA extracted from cells transfected with siScr or siNF90 were analyzed by small RNA-seq. Results are shown as log2 fold change versus –log10 p-value. (C) Total RNA extracted from cells described in (A) were analyzed by Taqman RT-qPCR as indicated. Results were normalized by those obtained for U6 abundance in the same samples. ND indicates “not detected”. Data represent mean ± SEM obtained from 3 independent experiments (\*\*\**P* < 0.001, independent Student’s *t* test).

The effect of NF90 on the abundance of miRNAs observed by miRNA profiling were validated by RT-qPCR analysis of selected miRNAs, miR-3173-3p, miR-186-5p, miR-1273c and miR-3189-3p, from biological triplicate samples. The results obtained confirmed the effects observed by miRNA profiling (Figure 1B, C). In addition, RNA was extracted from cells transfected with an independent non-targeting siRNA (Scr#2) and an NF90-targeting siRNA (NF90#2) that has been described previously (19) (Figure 1A, lower panel). Quantification of miRNAs 3173-3p, −186-5p, −1273c and - 3189-3p in biological triplicate samples (Figure 1C, lower panels) showed similar results to those obtained in Figure 1C upper panel, and also validated the results obtained by small RNA-seq. While we cannot exclude the possibility that a proportion of the small RNA-seq results could be due to off-target effects of the siRNAs, since only a single control and NF90-targeting siRNA were used, validation of a subset of the results using additional control and NF90-targeting siRNA suggests that the data are, to some extent, robust.

To evaluate whether the effect of NF90 on miRNA abundance might be cell type specific, we performed small RNA-seq in biological triplicate in HEK-293T cells transfected with control or NF90-targeting siRNA (Supplementary Figure S1A). Of 1917 annotated miRNA precursors, 1121, corresponding to 1647 mature miRNAs, were expressed in HEK-293T. Differential expression analysis (fold change ≥ 1.5 or ≤ 0.667; Adj*P-value* ≤ 0.05) revealed that 217 miRNA precursors, corresponding to 278 mature miRNAs, were upregulated following loss of NF90 while 77 precursors, corresponding to 84 mature miRNAs, were downregulated (Supplementary Figure S1B). Comparing upregulated miRNAs in the two cell types, we found 139 miRNAs that were upregulated in both cell lines after NF90 knock-down (Supplementary Figure S1C). This represents more than 65% of miRNAs upregulated in HepG2 and 64% of those upregulated in HEK-293T. Thus, NF90 appears to regulate a common subset of miRNAs.

### NF90 associates with a subset of pri-miRNAs

To determine which of the NF90-modulated miRNAs identified by miRNA profiling (Figure 1B) are direct targets of NF90, that is, pri-miRNAs that are bound by NF90, we took advantage of enhanced UV crosslinking followed by immunoprecipitation (eCLIP) dataset obtained in HepG2 cells (24). Analysis of HepG2 eCLIP data revealed 38 pri-miRNAs for which eCLIP peaks overlapped annotated pri-miRNA localizations +/-25 nt of flanking region (FC ≥ 1.5 and Bonferroni Adj*P* ≤ 0.05), as depicted in Figure 2A and Supplementary Table S5. Pri-miR-3173 and pri-miR-7-1 were among the 38 NF90-associated pri-miRNAs (Figure 2A, red dots).

**Figure 2.**
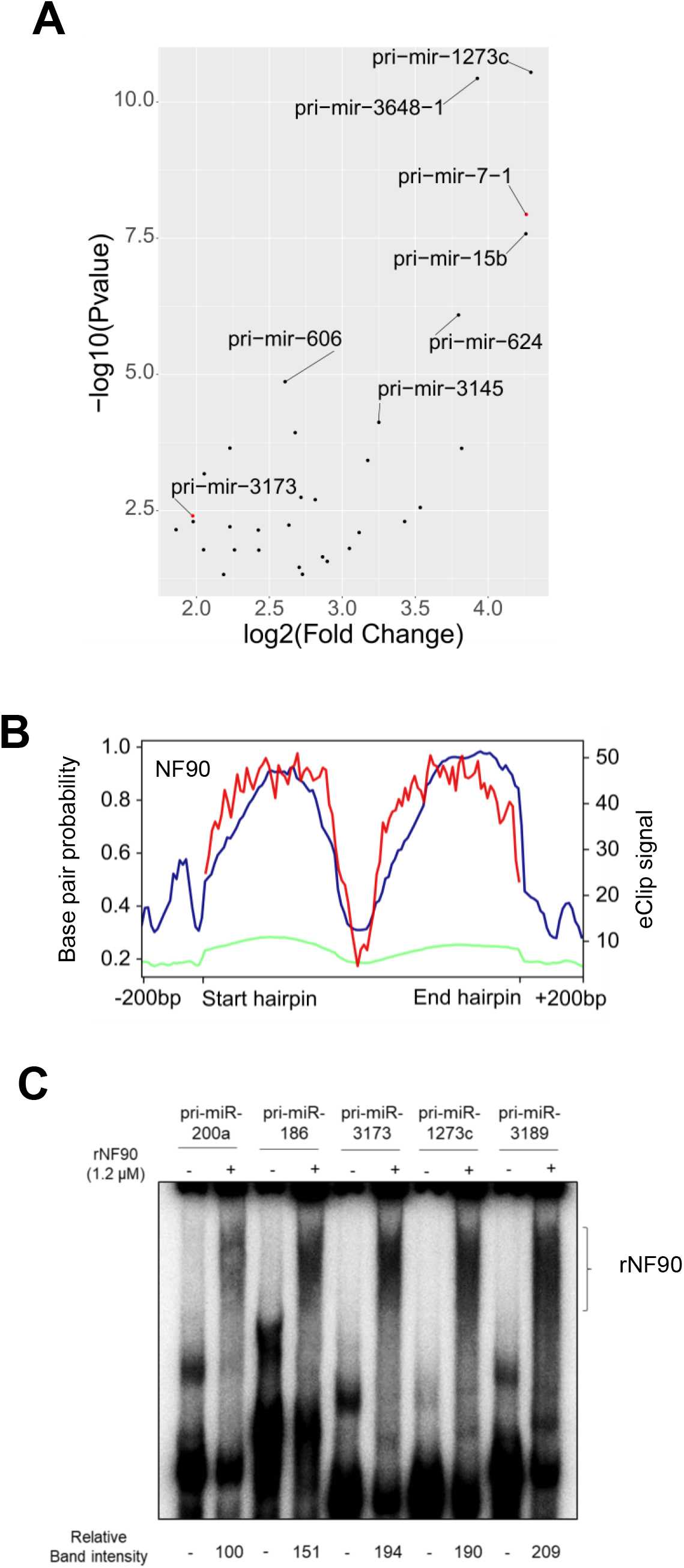
NF90 is associated with a subset of pri-miRNAs in HepG2 cells. (A) Dot plot representation of eCLIP data showing the 38 pri-miRNAs significantly associated with NF90 in HepG2 cells. Graph shows log2 fold change versus −log10 p-value. (B) Distribution of NF90 eCLIP reads along the region +/-200 bp of NF90-associated pri-miRNAs (blue) or all miRNAs (green) and base pair probability of NF90-associated hairpins (red). (C) RNA EMSA performed using recombinant NF90 was probed with radiolabelled pri-miRNAs as indicated. rNF90-pri-miRNA complexes are indicated on the figure.

We next analysed eCLIP read coverage across the pri-miRNA hairpin +/-200 bp for the 38 NF90-associated miRNAs compared to all pri-miRNAs (Figure 2B). As expected, analysis of all pri-miRNAs did not show significant read coverage for NF90 association. In contrast, NF90-associated miRNAs showed highest read coverage over the region having the strongest base pair probability and therefore likely corresponding to the double stranded pri-miRNA stem (Figure 2B). The region corresponding to the terminal loop, which has a low base pair probability, was not significantly bound by NF90. Interestingly, NF90 also appeared to bind to the pri-miRNA flanking region. Browser shots showing NF90 association with pri-miR7-1, pri-miR186 and pri-miR1273c by eCLIP are shown in Supplementary Figure S2.

To validate NF90 association with pri-miRNAs identified by eCLIP analysis (Figure 2A), we performed RNA EMSA using pri-miR-186, pri-miR-3173, pri-miR-1273c and pri-miR-3189 as radiolabelled probes together with recombinant NF90, as described previously for pri-miR7-1 and pri-miR-3173 (14,19). RNA EMSA confirmed NF90 association with pri-miR-186, pri-miR-3173, pri-miR-1273c and pri-miR-3189 (Figure 2C). Similarly, RNA EMSA confirmed that NF90 was not highly associated with pri-miR-200a, as indicated by eCLIP (Figure 2C).

Previous studies have indicated that NF90 may act as a competitor of Microprocessor for binding to pri-miRNAs (13-15,19). We therefore analysed eCLIP data for DGCR8 and Drosha performed in HepG2 cells (24). Association of DGCR8 was detected at 203 pri-miRNAs, while 147 pri-miRNAs were positive for Drosha binding (Figure 3A). Not surprisingly, there was a significant overlap between pri-miRNAs that were bound by both subunits of Microprocessor (Figure 3A). Indeed, 125 pri-miRNAs were associated with both factors, which represents approximately 60% and 85% of pri-miRNAs positive for DGCR8 and Drosha, respectively. Interestingly, only 10 pri-miRNAs bound by NF90 overlapped with those bound by either DGCR8 or Drosha, which represents approximately 24% overlap with DGCR8 and 13% overlap with Drosha (Figure 3A). This result indicates that NF90-associated pri-miRNAs are not highly associated with Microprocessor. Analysis of eCLIP reads showed association of DGCR8 with both apical and stem regions of pri-miRNAs (Figure 3B), as expected (27).

**Figure 3.**
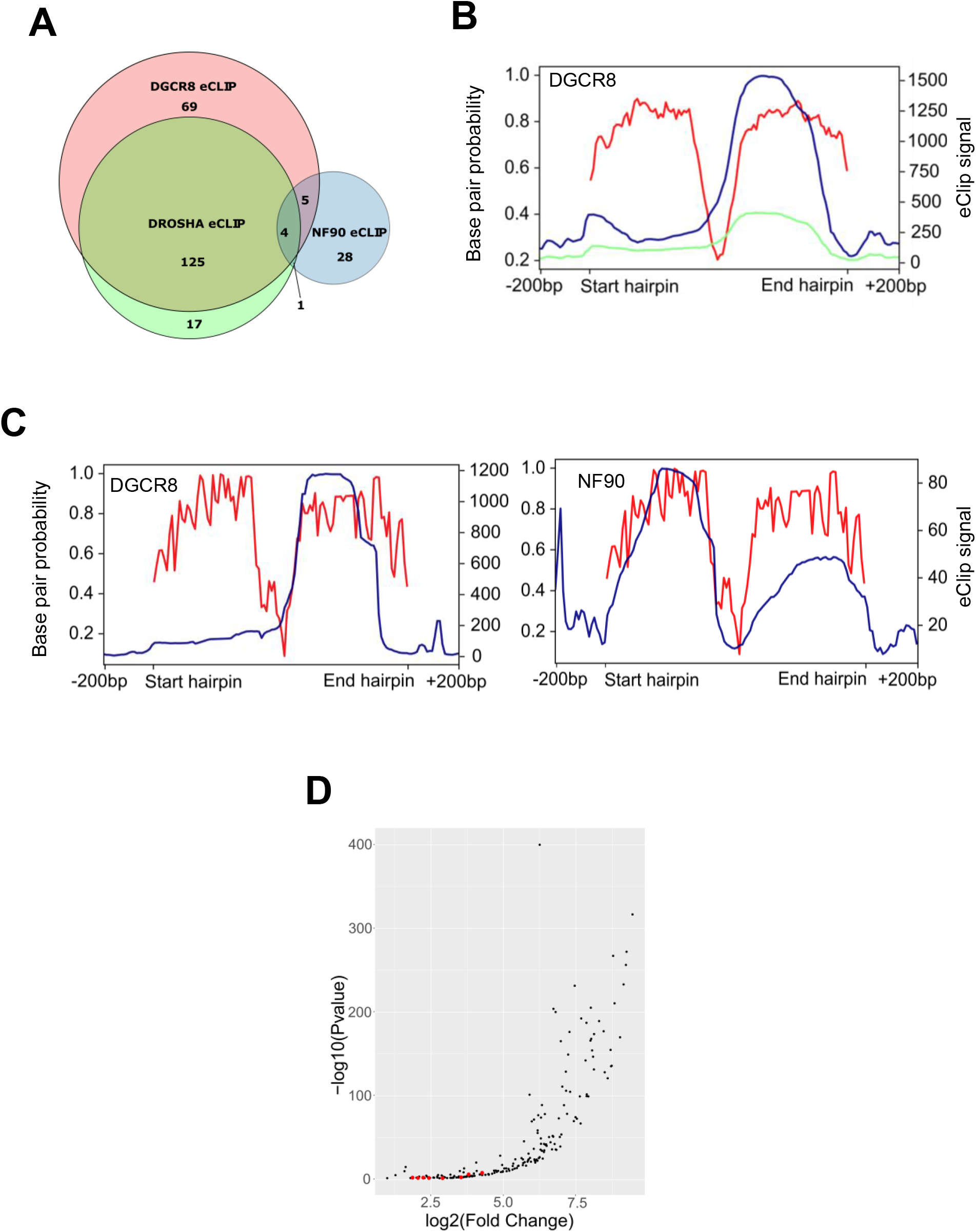
NF90-associated pri-miRNAs are poorly associated with Microprocessor. (A) Venn diagram showing the number of pri-miRNAs associated with DGCR8, Drosha or NF90 detected by eCLIP, as indicated. (B) Distribution of DGCR8 eCLIP reads along the region +/-200 bp of DGCR8-associated pri-miRNAs (blue) or all miRNAs (green) and base pair probability of DGCR8-associated hairpins (red). (C) Distribution of eCLIP reads along the region +/-200 bp of pri-miRNAs associated with both DGCR8 and NF90 (blue) and base pair probability of the hairpins (red). Left panel shows DGCR8 eCLIP reads, right panel shows NF90 eCLIP reads in blue. (D) Dot plot representation of eCLIP data showing 203 pri-miRNAs significantly associated with DGCR8 in HepG2 cells. Graph shows log2 fold change versus −log10 p-value. Red dots indicate pri-miRNAs that are also significantly associated with NF90.

We further analysed eCLIP read coverage over the pri-miRNAs that were found to be associated with both NF90 and DGCR8. While the profile for DGCR8 was similar to that for all DGCR8 positive pri-miRNAs (compare Figure 3C, left panel to Figure 3B), the profile for NF90 read coverage was somewhat different to that for all NF90-positive pri-miRNAs (compare Figure 3C, right panel, to Figure 2B). Interestingly, for pri-miRNAs that are bound by both DGCR8 and NF90, the profiles appear to be complementary (Figure 3C, compare left and right panels). Plot profiles of DROSHA and DGCR8 eCLIP data suggest that pri-miRNAs common with NF90 (shown with red dots) are not among the most enriched for Microprocessor binding (Figure 3D and Supplementary Figure S3).

### Pri-miRNAs that are bound and modulated by NF90 are highly stable

We next asked whether NF90 association with pri-miRNAs might affect their cropping by Microprocessor. If so, loss of NF90 would be predicted to increase the abundance of the mature miRNA products, as observed previously (14,15,19). MiRNA profiling revealed that of the 38 NF90-associated pri-miRNAs, 22 showed an increase in mature miRNA products, representing more than 57% of NF90-associated pri-miRNAs, while only 2 were decreased (Supplementary Tables S6 and S7). Thus, we identified a subset of 22 pri-miRNAs that are bound by NF90 and whose abundance is increased following loss of NF90, which we named ‘double-positive’ pri-miRNAs. Both pri-miR-7-1 and pri-miR-3173 were identified within the double positive subset. Thus, NF90 modulates the expression of most of its target pri-miRNAs.

Gene ontology of validated mRNA targets of double positive miRNAs revealed an implication particularly in cancer and infection by viruses, such as Epstein Barr Virus (EBV), hepatitis B virus (HBV), and human T lymphoma virus type 1 (HTLV1), as well as viral carcinogenesis (Supplementary Figure S4). This result is interesting given that NF90 translocates from the nucleus to the cytoplasm following viral infection of cells (28). Thus, viral infection could result in the coordinated processing of the NF90-modulated subset of pri-miRNAs, whose target mRNAs are implicated in viral replication.

We wondered whether pri-miRNAs that are associated with and modulated by NF90 might share a common characteristic that would make them targets for NF90 binding. A MEME search did not reveal a simple binding motif common to the 22 pri-miRNA sequences. Compared to all human pri-miRNAs, the subset of 22 double-positive pri-miRNAs did not show any significant difference in their overall length (mean=82.5 nt compared to 81.88 nt) or in the size of the terminal loop (mean=7.87 nt compared to 7.92 nt) (Figure 4A). In contrast, however, the minimal stretch containing a mismatch ≤1 nt was significantly longer for double-positive pri-miRNAs compared to all pri-miRNAs, with a mean of 27.68 nt for double-positive pri-miRNAs compared to 21.11 nt for all pri-miRNAs (Figure 4A). This analysis suggests that double-positive pri-miRNAs might be more stable, having a longer duplex and less bulges compared to all human pri-miRNAs. To further investigate this possibility, we compared the free energy of the 22 double-positive pri-miRNAs compared to all pri-miRNAs. The 22 double-positive pri-miRNAs had a lower free energy (mean=-42.26) compared to all pri-miRNAs (mean=-38.19), as shown in Figure 4B. Taken together, these data suggest that double positive pri-miRNAs are more stable and have less mismatches than all pri-miRNAs. Predicted folding of double-positive pri-miRNA sequences also revealed highly stable structures with very few bulges, compared to pri-miR-200a, which is not highly associated with NF90 (Supplementary Figure S5).

**Figure 4.**
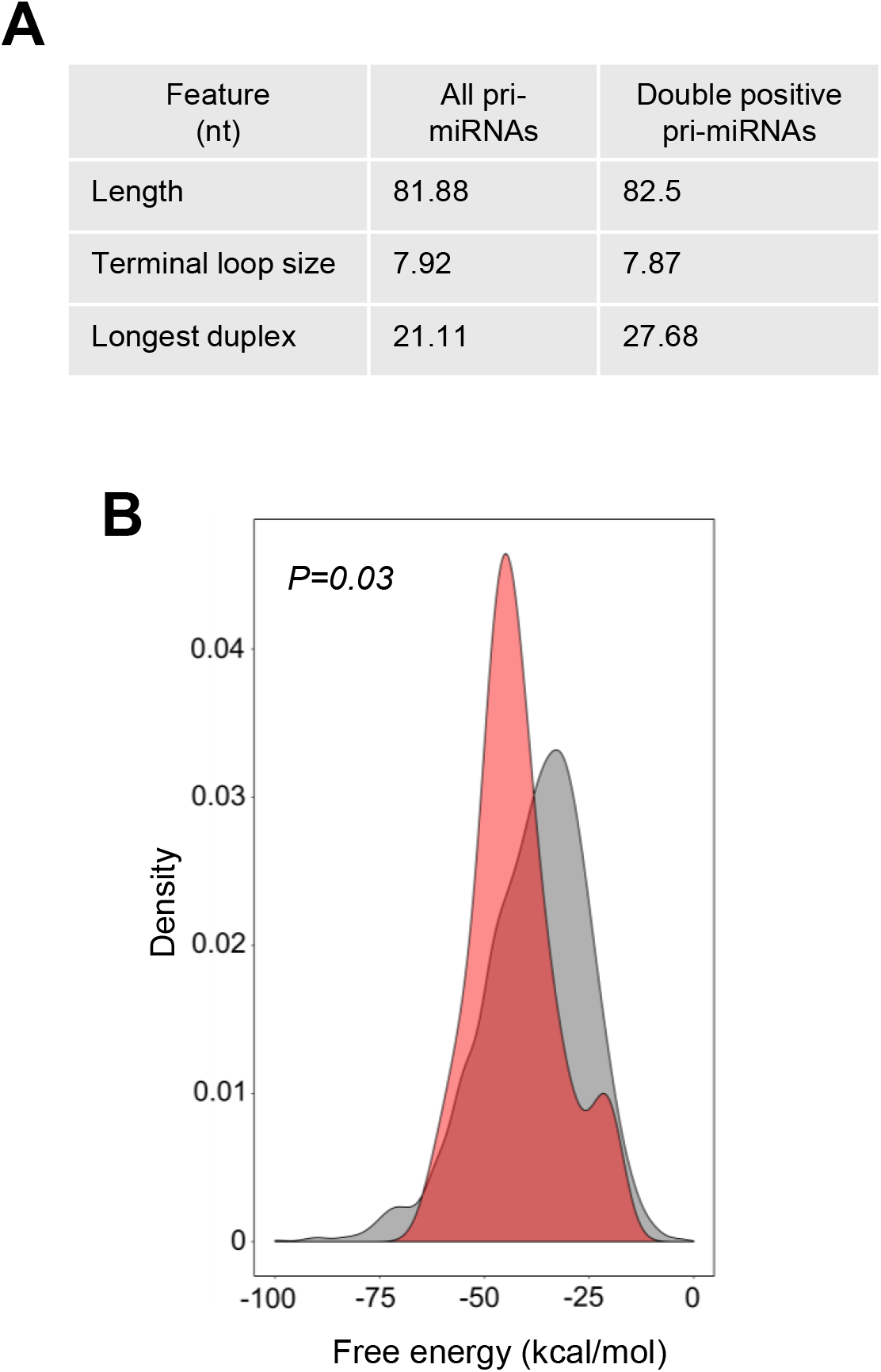
NF90 associates with a subset of highly stable pri-miRNAs. (A) Structural characteristics of all human pri-miRNAs and NF90 double positive pri-miRNAs. (B) Graph showing the free energy of all pri-miRNAs (grey) and NF90 double positive pri-miRNAs (red).

To test the idea that NF90 can bind to pri-miRNAs that have a stable structure with few bulges, we designed mutations within NF90-binding pri-miRNAs predicted to reduce stability and form bulge-like regions that might disrupt NF90 association. For each of the NF90-associated pri-miRNAs tested, we designed two mutant structures that would be less stable than wild-type structures. (Figure 5A). WT and mutated pri-miRNAs were tested for NF90 association by RNA EMSA. As shown in Figure 5B, mutation of pri-miR-3173 or pri-miR-186 to less stable structures diminished NF90 binding. On the other hand, mutation of pri-miR-200a to a more stable structure enhanced NF90 binding. These data suggest that NF90 shows a preference for association with stable pri-miRNA hairpin structures having few bulge regions.

**Figure 5.**
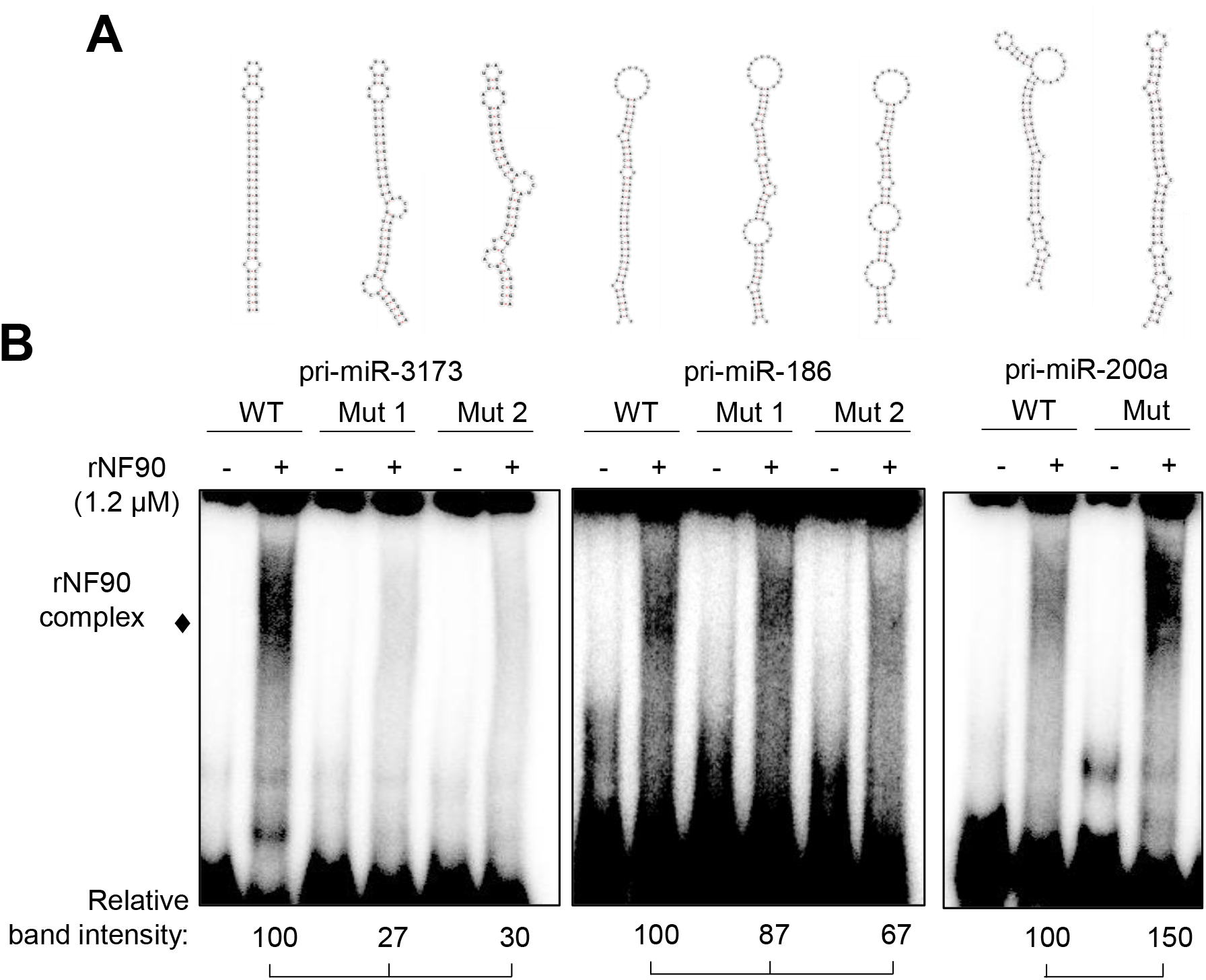
Modification of pri-miRNA structure alters NF90 binding. (A) Representations of wt or mutant pri-miRNAs sequences, as indicated. (B) RNA EMSA performed using recombinant NF90 and probed with radiolabelled pri-miRNAs as indicated. rNF90-pri-miRNA complexes are indicated on the figure. Relative band intensities (normalized to signal for wt) are shown below.

We then wondered whether pri-miRNAs whose mature products increased following loss of NF90, but were not considered eCLIP-positive using the applied cut-offs, might share the characteristics identified for double-positive pri-miRNAs. We therefore calculated the longest duplex length, allowing a mismatch of 1 nt, for the group of 181 upregulated but eCLIP negative pri-miRNAs, and 124 downregulated pri-miRNAs, as well as for those falling outside these groups (other) (Figure 6A). Interestingly, upregulated/eCLIP negative pri-miRNAs have a significantly longer duplex than all pri-miRNAs or other pri-miRNAs. Indeed, the duplex length is similar to that observed for the double positive group. In contrast, downregulated pri-miRNAs have a shorter duplex compared to all pri-miRNAs or other pri-miRNAs. We then calculated the mean free energy for the upregulated, eCLIP-negative group and the downregulated group of pri-miRNAs (Figure 6B). Similarly, when compared to all pri-miRNAs, the upregulated, eCLIP-negative group of pri-miRNAs had a significantly lower free energy. Free energy of the downregulated group was similar to that of all pri-miRNAs. In contrast, terminal loop size was comparable between the 2 groups; 7.86nt (downregulated group) compared with 8.64 nt (upregulated eCLIP-negative group). Of note, total pri-miRNA length was higher for the upregulated eCLIP-negative group (87.01 nt) compared to the downregulated group (77.79 nt). These analyses suggest that upregulated, eCLIP-negative pri-miRNAs share some characteristics with double-positive pri-miRNAs. It is feasible that some NF90-associated pri-miRNAs were not detected by eCLIP analysis or did not pass the selection criteria used to identify eCLIP-positive pri-miRNAs. To test this idea, we selected 2 pri-miRNAs, pri-miR-4755 and pri-miR-4766, from the upregulated, eCLIP-negative group whose structure corresponds to the defined criteria for NF90 association, that is, having low free energy and few mismatches (Supplementary Figure S6). NF90 binding to the pri-miRNAs was tested by RNA EMSA (Figure 6C). Indeed, both pri-miR-4755 and pri-miR-4766 were found to be associated with NF90.

**Figure 6.**
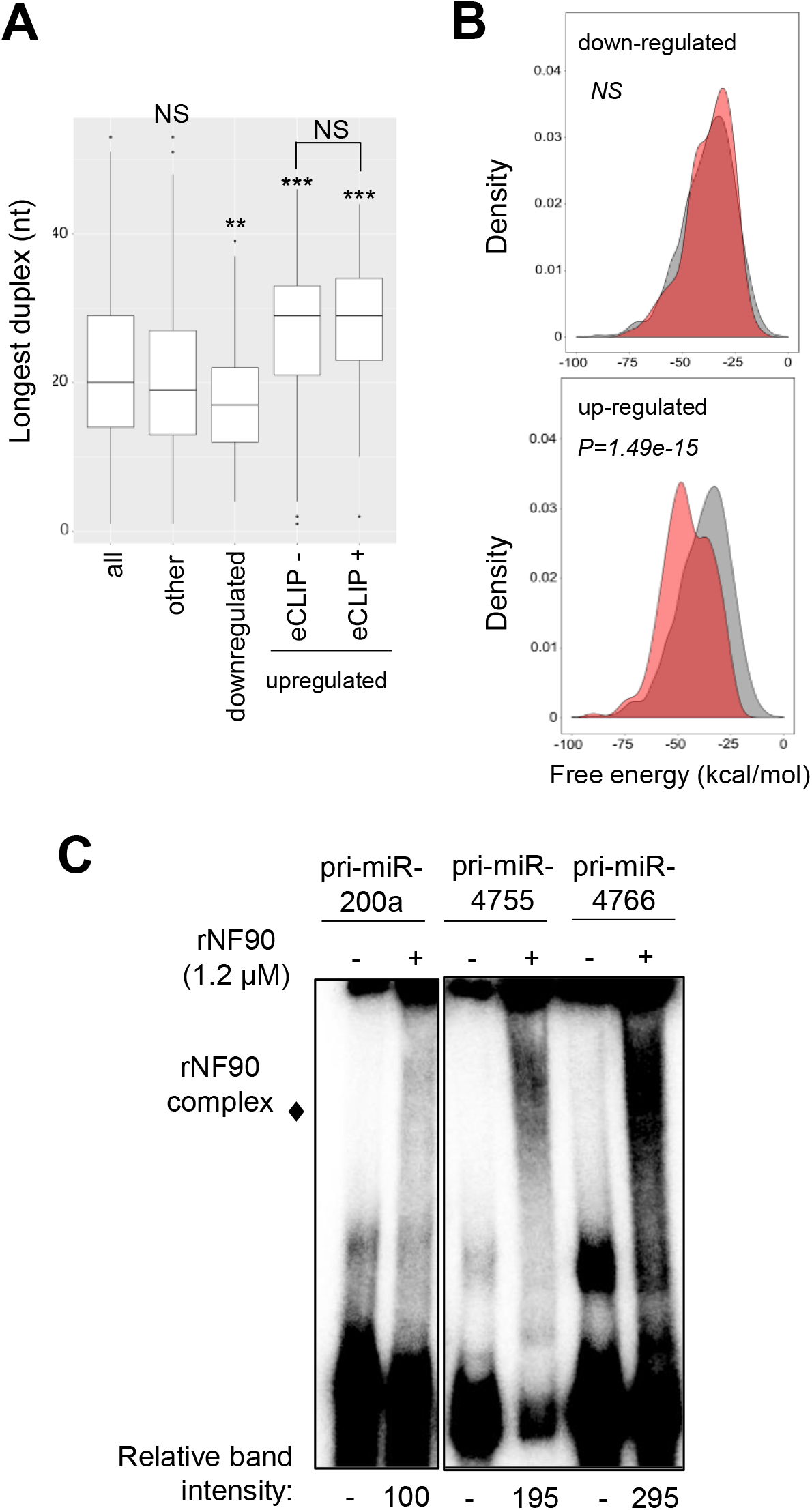
Pri-miRNAs whose mature products are upregulated following loss of NF90 share a similar structure. (A) Box plot representation of the longest duplex length of pri-miRNAs sorted into the indicated categories. (\**P* < 0.05, \*\*\**P* < 0.001, NS, not significant, Wilcoxon test). (B) Graphical representation of the free energy of pri-miRNAs whose mature products are downregulated or upregulated as indicated following loss of NF90 (red) compared to all pri-miRNAs (grey). (C) RNA EMSA performed using recombinant NF90 and probed with radiolabelled pri-miRNAs as indicated. rNF90-pri-miRNA complexes are indicated on the figure. Relative band intensities (normalized to pri-miR200a) are shown below.

### NF90 modulates the expression of a subset of genes hosting NF90-associated pri-miRNAs

Approximately 70% of human miRNAs are located in an intron of a host gene. Out of 22 double-positive pri-miRNAs, 20 are exclusively intronic. Two double-positive pri-miRNAs are found in either the 3’ UTR or an intron depending on transcript usage (Supplementary Table S6).

To determine whether loss of NF90 also affected the expression or splicing efficiency of the host genes, we performed RNA-seq in HepG2 cells transfected with control siRNA or siRNA targeting NF90. Loss of NF90 significantly diminished expression of 3 genes containing NF90-associated pri-miRNA; growth differentiation factor 15 (GDF15) hosting pri-miR-3189, 1-acylglycerol-3-phosphate O-acyltransferase 5 (AGPAT5) hosting pri-miR-4659a and zinc finger RAN-binding domain containing 2 (ZRANB2) hosting pri-miR-186 (Figure 7A). Furthermore, splicing efficiency of T-cell lymphoma invasion and metastasis 2 (TIAM2) hosting pri-miR-1273c was diminished (Figure 7B). In contrast, no significant effect was observed for NDUFS8, which hosts pri-miR-7113 and pri-miR-4691 that are not bound by NF90 and whose abundance are not modulated by NF90 (Figure 7B).

**Figure 7.**
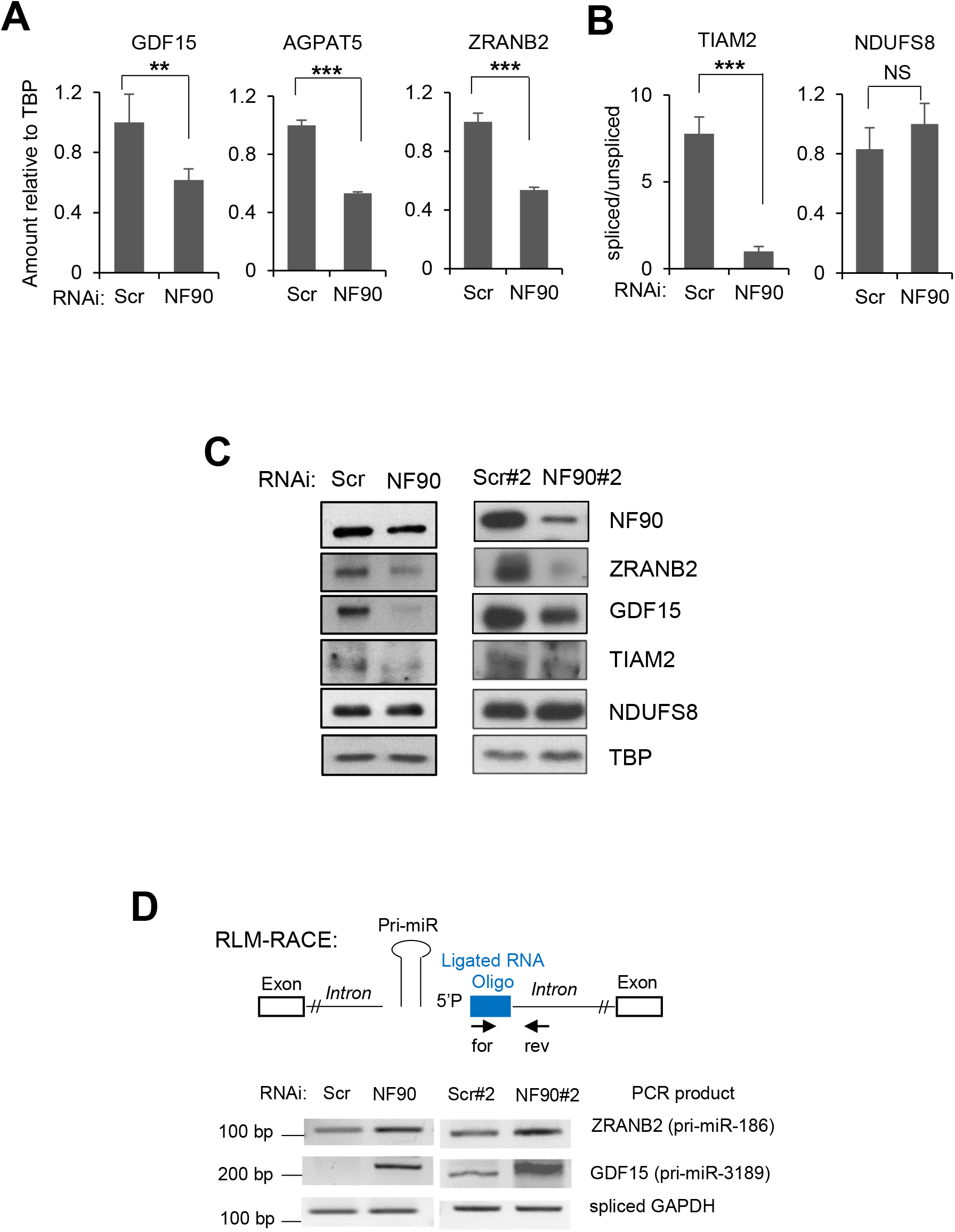
NF90 impacts expression of miRNA-hosting genes. (A) Extracts of HepG2 cells transfected with siRNA targeting NF90 or a non-targeting control (Scr) as indicated were analyzed by RNA-seq and DESeq2. Data represent mean ± SEM obtained from 3 independent samples (\*\**P* < 0.01, \*\*\**P* < 0.001, independent Student’s *t* test). (B) Exon-intron junction reads and exon-exon junction reads in samples described in A were counted using IRFinder software. The graph represents the mean ± SEM of the number of exon-exon junction reads divided by the number of exon-intron junction reads. (\*\*\**P* < 0.001, independent Student’s *t* test). **(C)** Extracts of HepG2 cells transfected with siRNA targeting NF90 (NF90, NF90#2) or a non-targeting control (Scr, Scr#2) as indicated were analyzed by immunoblot using the antibodies indicated. **(D)** NF90 modulates transcript cleavage at the region containing miRNA. Extracts of HepG2 cells transfected with siRNAs targeting NF90 (NF90, NF90#2) or non-targeting controls (Scr, Scr#2) as indicated were analyzed by modified 5’ RLM-RACE. Forward and reverse primers used, and the predicted sizes of the PCR products are indicated.

The expression of these genes was analysed by western blot of extracts obtained from HepG2 cells transfected with control (Scr and Scr#2) and NF90-targeting (NF90 and NF90#2) siRNAs. All genes tested showed diminished expression upon loss of NF90, except NDUFS8 that showed no significant difference in expression (Figure 7C). Thus, NF90 modulates the expression of certain pri-miRNA host genes, including TIAM2, a known oncogene and metastasis factor in HCC (29,30).

Finally, to determine whether loss of gene expression correlated with increased pri-miRNA cropping following knock down of NF90, we performed modified RLM-5’ RACE as described previously (19), using extracts of cells transfected with control (Scr and Scr#2) and NF90-targeting (NF90 and NF90#2) siRNAs. Indeed, RLM RACE analysis showed enhanced cleavage of the intronic region of ZRANB2 hosting pri-miR-186 and GDF15 hosting pri-miR-3189 in extracts of NF90 knock down cells compared to controls (Figure 7D). This analysis indicates that loss of NF90 enhances transcript cleavage in the vicinity of the hosted pri-miRNA.

## DISCUSSION

We and others have previously shown that NF90 can modulate the processing of certain miRNA precursors (14,15,19). However, it was unclear how widespread the impact of NF90 might be on human miRNA biogenesis. Here, we have used genome-wide approaches to address the effect of NF90 on the miRNA pool in HepG2 HCC cells. Our data indicate that NF90 modulates the processing of a specific subset of miRNA precursors. NF90 is associated with at least 38 human pri-miRNAs, as indicated by analysis of eCLIP data obtained by Nussbacher and Yeo (24). Of these, 22 showed increased abundance of mature miRNA products following knock-down of NF90. Thus, association of NF90 with a pri-miRNA is likely to influence its fate. Most NF90-associated pri-miRNAs did not overlap with those bound by either DGCR8 or Drosha. Interestingly, for those pri-miRNAs that were bound by both NF90 and DGCR8, the binding profiles of the two factors were largely complementary. Furthermore, while the binding profile of DGCR8 was not noticeably different for this group compared to all pri-miRNAs bound by DGCR8, the binding profile of NF90 differed somewhat for this group compared to all pri-miRNAs bound by NF90. This could suggest that NF90 and DGCR8 might bind simultaneously to the pri-miRNA, and that the binding of DGCR8 may alter the binding mode of NF90 for such pri-miRNAs.

Since NF90 is a highly abundant and ubiquitously expressed protein, it might be expected that NF90-associated pri-miRNAs would be poorly processed in most cells. Indeed, the mature miRNA products of NF90 bound pri-miRNAs are very poorly expressed, or not expressed at all in control cells. They become readily detectable only upon loss of NF90. An exception is pri-miR-7-1, although interestingly, this miRNA shows tissue specific expression, being highly expressed only in brain and pancreas (31).

Our data suggests that NF90-moduated pri-miRNAs share a common structure that might facilitate NF90 association with the stem region. This finding is consistent with a previous report showing structure-based recognition of adenovirus-expressed VA1 RNA by NF90 (32). Extensive mutational analysis of VA1 association with NF90 showed no specificity for nucleotide sequence but rather the requirement for a minihelix structure within the stem region. The pri-miRNAs identified in this study also exhibit a minihelix-like structure that appears to be necessary for NF90 binding. Indeed, RNA EMSA showed that NF90 association with pri-miR-3173 and pri-miR-186 could be diminished by introducing destabilizing mutations, while NF90 association could be acquired by increasing the stability of the stem region, as for pri-miR-200a.

Interestingly, our data predict that the subset of NF90-associated pri-miRNAs may extend beyond those detected by eCLIP analysis. Using the characteristics determined from the eCLIP-positive, upregulated pri-miRNA group, that is duplex length and free energy, we found that pri-miRNAs whose mature products were upregulated following loss of NF90 but were not positive by eCLIP analysis shared the same characteristics as the double positive group. The length of the duplex region and the free energy of the structure was comparable to that of double positive pri-miRNAs. RNA EMSA confirmed the predicted association with NF90 for two of these pri-miRNAs. Interestingly, both groups were significantly different to all pri-miRNAs or those that are unaffected by NF90 (other). Thus, it appears that the high specificity of eCLIP revealed a subset of pri-miRNAs that share a common structure. When this information was used to interrogate the group of pri-miRNAs who share the same biological response to loss of NF90, that is, upregulation of their mature products, we observed that both groups share the same characteristics. We predict that a certain number of the upregulated group likely do bind to NF90 but may escape detection by eCLIP. For example, as noted above, many of the pri-miRNAs are expressed at extremely low levels in control cells, which could make their association with NF90 difficult to detect.

Interestingly, pri-miR-7-1 processing has been shown to be influenced by another RBP, HuR, which recruits MSI2 to the terminal loop. Binding of HuR/MSI2 was found to stabilize the stem region and led to diminished processing by Microprocessor (33). It would be interesting to determine whether binding of HuR/MSI2 to pri-miR-7-1 might facilitate NF90 binding to the stem region, and compete with Microprocessor. Similarly, it would be interesting to determine whether HuR/MSI2 can bind the terminal loop of other NF90-modulated pri-miRNAs in addition to pri-miR-7-1. NF90 may cooperate with other RBPs, such as HuR/MSI2 to control the processing of a subset of pri-miRNAs.

Another feature that the subset of NF90-modulated pri-miRNAs share is their restriction to human or primate lineages. Again, pri-miR-7-1 is an exception, being highly conserved throughout evolution. Thus, given that the subset of NF90-modulated pri-miRNAs are young and almost perfect hairpins, it is tempting to speculate that this group may have originated through recent insertion of repeat elements in the genome. Of note, NF90 was found to associate with and modulate the abundance of several members of the primate-specific miR-548 family, which are derived from Made1 transposable elements. Made1 elements are short miniature inverted-repeat transposable elements (MITEs) consisting of two 37 nt terminal inverted repeats that flank 6 bp of internal sequence. Thus, Made1 elements are nearly perfect palindromes, and when expressed as RNA they form highly stable hairpin loops. There are more than 3,500 putative miR-548 target genes in humans. Analysis of their expression profiles and functional affinities suggests cancer-related regulatory roles for miR-548 family members (34). It is possible that NF90 may be required to diminish their processing under control conditions.

Interestingly, GO analysis of validated mRNA targets of the mature miRNAs showed significant enrichment for infection by viruses such as Epstein Barr Virus (EBV), hepatitis B virus (HBV) and human T lymphoma virus type 1 (HTLV1) and in viral carcinogenesis. Indeed, viral infection of cells induces translocation of NF90 from the nucleus to the cytoplasm (28). Thus, it is conceivable that pathological conditions such as viral infection could result in the coordinated processing of the NF90-modulated subset of pri-miRNAs, which target mRNAs important for viral replication.

Finally, transcriptomic analysis showed that association of NF90 with pri-miRNAs may modulate expression of certain host genes, as described previously (19). Among the modulated host transcripts hosting pri-miRNAs, two are noteworthy. The expression of TIAM2, hosting pri-miR-1273C, is down-regulated upon loss of NF90. TIAM2 is a known oncogene and metastasis factor in HCC (29,30). Levels of NF90 are elevated in HCC (14,20) and it would be interesting to determine whether NF90-dependent modulation of TIAM2 might contribute to pathogenesis. Loss of NF90 also diminished expression of growth differentiation factor 15 (GDF15), hosting pri-miR-3189. GDF15 is expressed and secreted by a limited number of tissues, including liver. When complexed with its receptor, GFRAL, in brain and CNS, GDF15 supresses appetite (see (35) for review). Cancer patients express high circulating levels of GDF15, which contributes to anorexia/cachexia. On the other hand, enhancement of GDF15 expression is a promising therapeutic strategy in the treatment of obesity. It would be interesting to determine whether high levels of NF90 in HCC may have a role in promoting expression of GDF15 from liver cells in cancer patients.

In summary, we have identified a subset of human pri-miRNAs that are bound by NF90. Analysis indicates that this subset shares a similar structure that appears to be favourable for NF90 binding. These data extend our knowledge of how processing of pri-miRNAs can be modulated by RBPs. This may be beneficial for understanding perturbations of miRNA levels in pathological conditions and could also open up novel treatment strategies using nanotherapeutics.

## ACCESSION NUMBERS

Small RNA-seq and RNA-seq data have been deposited at GEO (GSE132341).

## ACKNOWLEDGEMENT

We wish to thank Catherine Dargemont and Xavier Contreras for critical reading of the manuscript, and the Gene Regulation lab and Hervé Seitz for helpful discussions.

