## Supplementary material for "NF90 Modulates Processing of a Subset of Human Pri-miRNAs": Suppl Figs and Tables

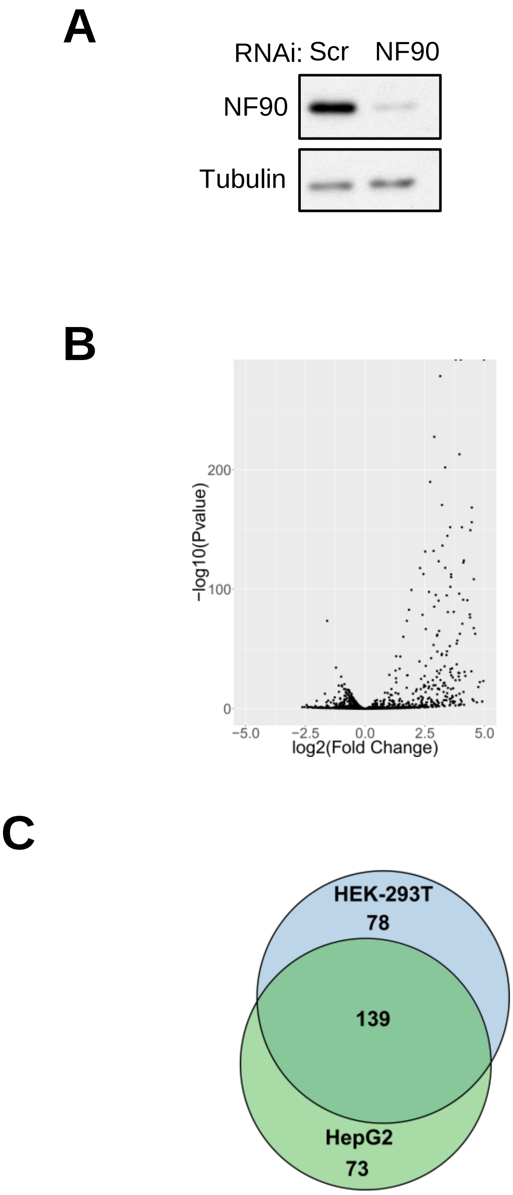

**Figure S1.** NF90 Modulates the Expression Level of miRNAs in HEK-293T cells. **(A)** Extracts of HEK-293T cells transfected with siRNA targeting NF90 or a non-targeting control (Scr) were analyzed by Western blot using the indicated antibodies. **(B)** Samples described in A were analyzed by small RNA-seq. Results are shown as log2 fold change versus –log10 p-value. **(C)** Venn diagram representing the number of miRNAs upregulated following knock-down of NF90 in HepG2 versus HEK-293T cells.

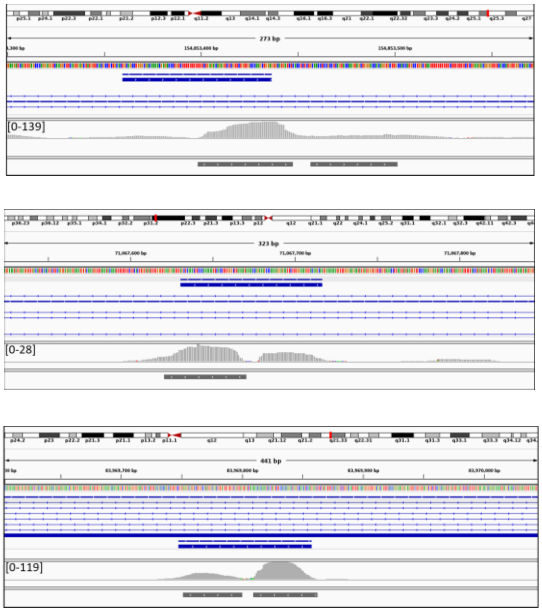

pri-miR-1273C
TIAM2

pri-miR-186
ZRANB2

HNRNPK
pri-miR-7-1

**Figure S2.** Browser shots of NF90 eCLIP read coverage over the pri-miRNAs indicated on the figure. Blue lines represent host gene showing localization of the pri-miRNA. eCLIP reads are shown in grey and locations of eCLIP peaks are shown as dark grey bars. The arrows indicate the strand from which the reads originated.

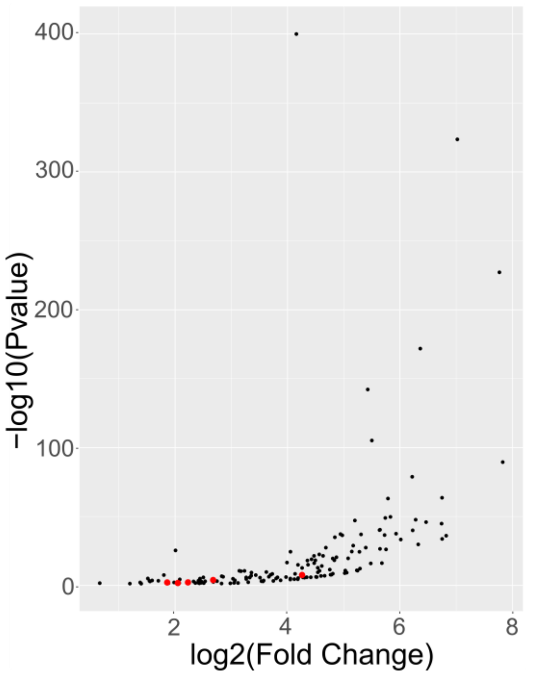

**Figure S3.** DotPlot of Drosha-associated pri-miRNAs, determined by eCLIP analysis. Red dots indicate the position of pri-miRNAs that are also positive for association with NF90.

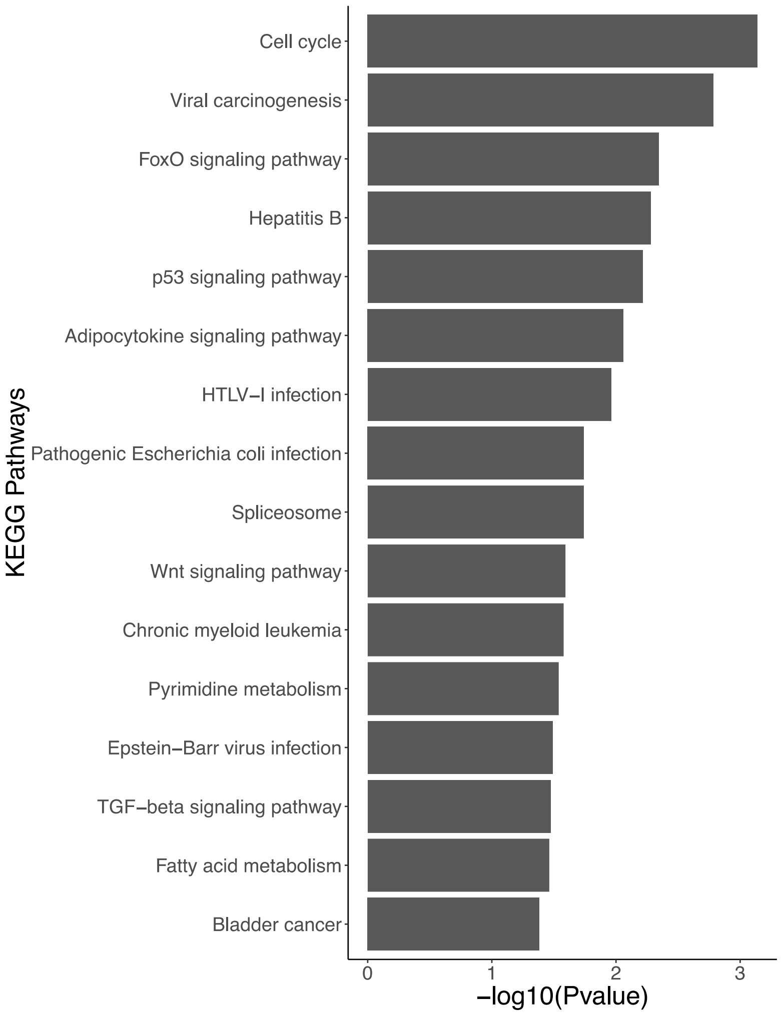

**Figure S4.** NF90 double positive pri-miRNAs target genes involved in viral infection and cancer. Gene ontology of validated targets of NF90-bound and upregulated miRNAs.

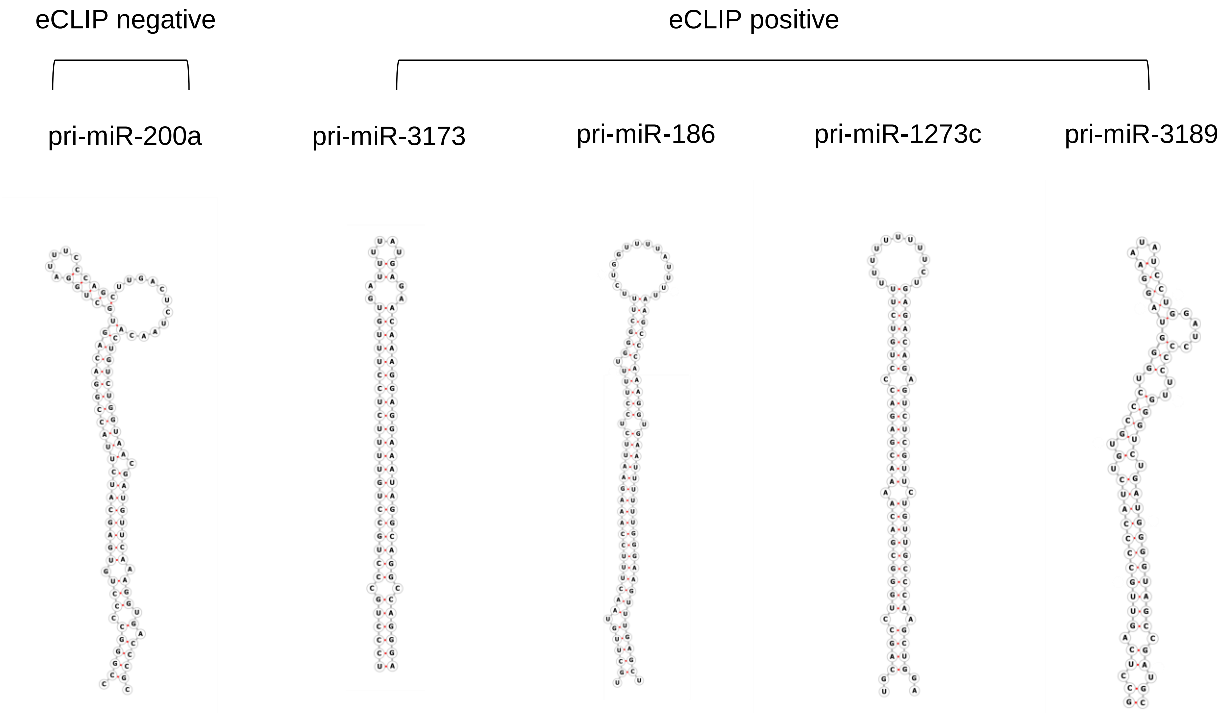

**Figure S5.** NF90-associated pri-miRNAs are highly stable. Predicted folding of pri-miRNAs that are significantly associated or not with NF90 as indicated. RNA structures were predicted using FORNA.

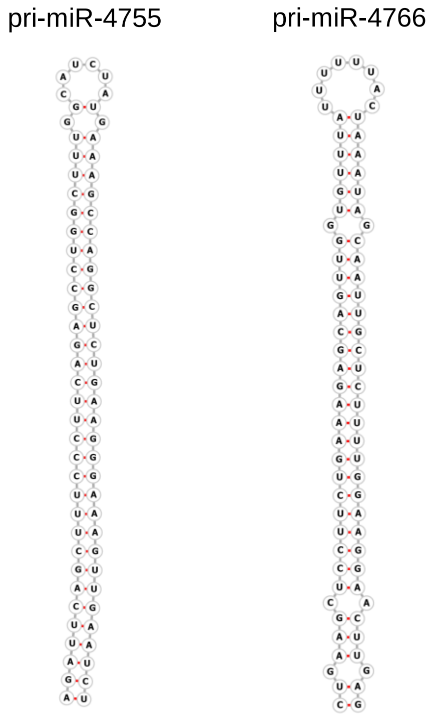

**Figure S6.** NF90-modulated pri-miRNAs are highly stable. Predicted folding of examples of pri-miRNAs whose mature products are upregulated following loss of NF90. RNA structures were predicted using FORNA.

**Supplementary Table S1.** Double stranded siRNAs used in this study.

| siRNA | Sequence (5' to 3') |
|---|---|
| Scr | gcgcgcuuuguaggauucg(dTdT) |
| Scr#2 | ucugcaagguuaggcgucu(dTdT) |
| NF90 | ccaaggaacucuaucacaa(dTdT) |
| NF90#2 | gaguugaaguauugauaac(dTdT) |

**Supplementary Table S2.** Primary antibodies used in this study.

| Antibody | Reference | Supplier |
|---|---|---|
| NF90 | A303-651A | Bethyl Laboratories |
| GDF-15 | sc-377195 | SCBT |
| ZRANB2 | sc-514200 | SCBT |
| TIAM2 | sc-514090 | SCBT |
| NDUFS8 | sc-515527 | SCBT |
| TUBULIN | DM1A clone, T6199 | Sigma-Aldrich |
| TBP | sc-421 | SCBT |

**Supplementary Table S3.** Primers used in this study.

| Primer | Forward (5' to 3') | Reverse (5' to 3') |
|---|---|---|
| Spiced GAPDH | cac atc gct cag aca cca t | gag gtc aat gaa ggg gtc at |
| U6 | ctc gct tcg gca gca cat ata c | gga acg ctt cac gaa ttt gcg tg |
| pri-miR-1273C | ctt ggg aag ctg agg tag gc | act tgg tac tga ggc gga gg |
| pri-miR-186 | aca gaa cac cca tca tat tc | gtt gac att cac atg ctt c |
| pri-miR-200a | ctg gct gct cac cgc tcc | gat gtg cct cgg tgg tgt cc |
| pri-miR-3173 | cat tgg agg tct agg gct ta | gtt ctt cct cgg cac aag |
| pri-miR-3189 | agc agc ccc cat atc taa tc | ctg gca tcc ctg tac ctc |
| pri-miR-4755 | aga gat gag gaa ggt tat ggc t | tgg ccc aaa cct cat aga c |
| pri-miR-4766 | ccc ttc tac ctt tct gaa gct c | cac aca ggt ggc act caa c |
| 5'RLM-RACE pri-miR-186 (outer) | gct gat ggc gat gaa tga aca ctg (adapter) | aaa cca ggt ata tgg cac agc aac |
| 5'RLM-RACE pri-miR-186 (inner) | cgc gga tcc gaa cac tgc gtt tgc tgg ctt tga tg (adapter) | tgt tga cat tca cat gct tca ggt |
| 5'RLM-RACE pri-miR-3189 (outer) | gct gat ggc gat gaa tga aca ctg (adapter) | acc aca ccc cca ttg ttt ctct |
| 5'RLM-RACE pri-miR-3189 (inner) | cgc gga tcc gaa cac tgc gtt tgc tgg ctt tga tg (adapter) | acc aca ccc cca ttg ttt ctct |

**Supplementary Table S4.** Wild-type and mutant pri-miRNAs sequences used for RNA EMSA. The pri-miRNA sequence is shown in red and flanking sequence is shown in black.

| | pri-miR-200a | pri-miR-3173 | pri-miR-186 |
|---|---|---|---|
| Wild-type | ctggctgctcaccgctccggttcttccctgggcttccacagcagcccctgcctgcctggcgggaccccacgtccctcccgggcccctgtgagcatcttaccggacagtgctggatttcccagcttgactctaacactgtctggtaacgatgttcaaaggtgacccgccgctcgccggggacaccaccgaggcacatc | cattggaggtctagggcttattttccagatagaattgagtctttgttggtcttgggccagcttccctgccctgcctgttttctcctttgtgattttatgagaacaaaggaggaaataggcaggccagggaaacgatctctctccctctcttgtgccgaggaagaact | acagaacacccatcatattcttcccaaacattttttcattgcttgtaactttccaaagaattctccttttgggctttctggttttattttaagcccaaaggtgaatttttgggaagtttgagctaaattccttcaaccaaaatatacaagtgaagaaaaaaaatttgtatttaaacatttgcacatttacttctacctgaagcatgtgaatgtcaac |
| Mutant #1 | ctggctgctcaccgctccggttcttccctgggcttccacagcagcccctgcctgcctggcgggaccccacgtccctcccgggcccctgtgagcatcttaccggacagtgctggatttcccagcctgtctggtaacgatgttcaaaggtgacccgccgctcgccggggacaccaccgaggcacatc | cattggaggtctagggcttattttccagatagaattgagtctttgttggtcttgggccagcttccctgcgacgcctgcctgttttctcctttgtgattttatgagaacaaaggaggaaagcgctaggcaggccagggaaacgatctctctccctctcttgtgccgaggaagaact | acagaacacccatcatattcttcccaaacattttttcattgcttgtaactttccaactaaagaattctccttttgggctttctggttttattttaagcccacctctaaggtgaatttttgggaagtttgagctaaattccttcaaccaaaatatacaagtgaagaaaaaaaaatttgtatttaaacatttgcacatttacttctacctgaagcatgtgaatgtcaac |
| Mutant #2 | | cattggaggtctagggcttattttccagatagaattgagtctttgttggtcttgggccagcttccctgcaagtcctgttttctcctttgtgattttatgagaacaaaggagaccctaggcaggccagggaaacgatctctctccctctcttgtgccgaggaagaact | acagaacacccatcatattcttcccaaacattttttcattgcttgtccagttccaaagaattctccttttgggctttctggttttattttaagcccaaaggtgaaccacgtgggaagtttgagctaaattccttcaaccaaaatatacaagtgaagaaaaaaaaatttgtatttaaacatttgcacatttacttctacctgaagcatgtgaatgtcaac |

**Supplementary Table S5.** NF90-associated pri-miRNAs, as determined by eCLIP analysis.

| | | |
|---|---|---|
| hsa-mir-1273c | hsa-mir-4485 | hsa-mir-548d-1 |
| hsa-mir-1290 | hsa-mir-4635 | hsa-mir-548u |
| hsa-mir-15b | hsa-mir-4659a | hsa-mir-548v |
| hsa-mir-186 | hsa-mir-4687 | hsa-mir-5581 |
| hsa-mir-1914 | hsa-mir-4712 | hsa-mir-570 |
| hsa-mir-3140 | hsa-mir-4714 | hsa-mir-578 |
| hsa-mir-3145 | hsa-mir-4730 | hsa-mir-579 |
| hsa-mir-3173 | hsa-mir-4762 | hsa-mir-606 |
| hsa-mir-3189 | hsa-mir-4775 | hsa-mir-624 |
| hsa-mir-3646 | hsa-mir-4779 | hsa-mir-6751 |
| hsa-mir-3648-1 | hsa-mir-4782 | hsa-mir-6839 |
| hsa-mir-3680-1 | hsa-mir-548aq | hsa-mir-7-1 |
| hsa-mir-3939 | hsa-mir-548ar | |

**Supplementary Table S6.** 'Double positive' miRNAs whose abundance increased following loss of NF90 and that were positive for NF90 association by eCLIP, and their host gene.

| miRNA | Small RNA-seq | | Host Gene |
|---|---|---|---|
| | Fold Change (log2) | p value (-Log10) | |
| hsa-mir-1273c | 3.55 | 33.94 | TIAM2 |
| hsa-mir-186 | 1.01 | 4.55 | ZRANB2 |
| hsa-mir-3140 | 3.77 | 37.16 | FBXW7 |
| hsa-mir-3145 | 2.56 | 9.78 | NHSL1 |
| hsa-mir-3173 | 3.03 | 25.36 | DICER1 |
| hsa-mir-3189 | 2.16 | 19.36 | GDF15 |
| hsa-mir-3646 | 3.76 | 4.88 | HNF4A |
| hsa-mir-3939 | 1.27 | 7.59 | RP1-167A14.2 |
| hsa-mir-4659a | 2.21 | 10.93 | AGPAT5 |
| hsa-mir-4714 | 3.36 | 23.31 | IGF1R |
| hsa-mir-4762 | 2.14 | 5.66 | ATXN10 |
| hsa-mir-4775 | 1.37 | 3.72 | CCNYL1 |
| hsa-mir-4779 | 4.03 | 10.9 | IMMT |
| hsa-mir-4782 | 3.94 | 3.87 | SLC35F5 |
| hsa-mir-548ar | 3.24 | 6.79 | CDC16 |
| hsa-mir-548u | 2.24 | 2.38 | PRIM2 |
| hsa-mir-548v | 2.01 | 7.81 | MTUS1 |
| hsa-mir-5581 | 2.69 | 19.23 | MEAF6 |
| hsa-mir-578 | 2.17 | 3.18 | CPE |
| hsa-mir-579 | 4 | 73.29 | ZFR |
| hsa-mir-624 | 2.68 | 25.49 | STRN3 |
| hsa-mir-7-1 | 0.99 | 5.76 | HNRNPK |

**Supplementary Table S7.** MiRNAs downregulated in abundance following loss of NF90 and that are associated with NF90 by eCLIP.

| | | | |
|---|---|---|---|
| miRNA | Fold Change (log2) | p value (-Log10) | Host Gene |
| hsa-mir-1914 | -0.86 | 2.88 | UCKL1 |
| hsa-mir-6751 | -1.52 | 3.73 | SYVN1 |
